# Developmental plasticity in male courtship in *Bicyclus anynana* butterflies is driven by hormone regulation of the *yellow* gene

**DOI:** 10.1101/2021.06.13.448264

**Authors:** Heidi Connahs, Eunice Jingmei Tan, Yi Ting Ter, Emilie Dion, Yuji Matsuoka, Ashley Bear, Antónia Monteiro

## Abstract

The organizational role for hormones in the regulation of sexual behavior is currently poorly explored. Previous work showed that seasonal variation in levels of the steroid hormone 20-hydroxyecdysone (20E) during pupal development regulates plasticity in male courtship behavior in *Bicyclus anynana* butterflies. Wet season (WS) males, reared at high temperature, have high levels of 20-hydroxyecdysone (20E) during pupation and become active courters. Dry season (DS) males, reared at low temperatures, have lower levels of 20E and lower courtship rates. Rescue of WS courtship rates can be achieved via injection of 20E into DS male pupae, but it is still unknown whether 20E alters gene expression in the pupal brain, and if so, the identity of those targets. Using transcriptomics, qPCR, and behavioral assays with a transgenic knockout, we show that higher expression levels of the *yellow* gene in DS male pupal brains, relative to WS brains, represses courtship in DS males. Furthermore, injecting DS males with 20E downregulates *yellow* to WS levels 4 hours post-injection, revealing a hormone sensitive window that determines courtship behavior. These findings are in striking contrast to *Drosophila*, where *yellow* is required for active male courtship behavior. We conclude that 20E plays an organizational role during pupal brain development by regulating the expression of *yellow,* which is a repressor of the neural circuity for male courtship behavior in *B. anynana*. This work shows that similar to vertebrates, hormones can also play an organizational role in insect brains, leading to permanent changes in adult sexual behavior.

**Significance Statement:** Behavioral plasticity in adult insects is known to be regulated by hormones, which activate neural circuits in response to environmental cues. Here, we show that hormones can also regulate adult behavioral plasticity by altering gene expression during brain development, adjusting the insect’s behavior to predictable seasonal environmental variation. We show that seasonal changes in the hormone 20E alters expression of the *yellow* gene in the developing pupal brain of *Bicyclus anynana* butterflies, which leads to differences in male courtship behavior between the dry and wet seasonal forms. This work provides one of the first examples of the organizational role of hormones in altering gene expression and adult sexual behavior in the developing insect brain.

## Introduction

Behavioral plasticity is essential for animals to adapt to environmental variation and it is often triggered by hormonal changes that organize or activate neural circuits in the brain (1–3). Seasonal changes in temperature and photoperiod can serve as important cues that alter hormone levels and sexual behavior in a wide range of vertebrate and invertebrate taxa (4). Precisely how hormone signaling influences sexual behavior in most animals however, is not well known (5). In vertebrates, hormones are considered to play both a brain organizational role, during development, as well as a behavioral activational role in adults, compared to just an adult activational role in insects (6, 7). This conceptual framework describes whether hormones permanently organize neural circuitry during early critical periods that later influence adult behavior, or whether they modulate behavior by activating existing neural circuits in response to external cues (7, 8).

Sexual behavior in insects has traditionally been viewed as a consequence of cell-autonomous processes taking place during brain development, and involving sex determination genes (6) such as *fruitless* and *doublesex* (9). The role of insect hormones is typically described as playing an activational role, allowing rapid and reversible behavioral changes, such as activating neural circuits that regulate pheromone communication or sexual receptivity (10–13). However, hormones have also been proposed to play an organizational role in insects, for instance in the regulation of behavioral polyphenisms in honeybees and locusts (7) and sexual maturity in *Drosophila* (14). Yet, no evidence is available for the organizational role of steroid hormones in driving sexual behaviors in adult insects, similar to the role of steroid hormones in vertebrate sexual differentiation, where exposure to different hormone levels during ontogeny leads to discrete, fixed differences in neural development and sexual behaviors (7).

For insects living in seasonal environments, hormones could play an organizational role earlier in development to ensure that sexual behavior is optimized for particular environmental conditions that will be prevalent upon adult emergence (15). An example of a species where such a brain organizational role may be happening is the African seasonal polyphenic butterfly, *Bicyclus anynana*. This species shows an interesting sex role reversal between seasonal forms that develop at different temperatures, and where temperature cues in the arrival of different seasons. In particular, wet season (WS) males, reared at high temperatures, play the active courting role, while DS males, reared at low temperatures, do much less courtship and become, instead, the choosy sex (16). The adaptive reason driving courtship plasticity in males is associated with increased reproductive costs for DS males, which provide females with beneficial spermatophores (16). Provision of this spermatophore ultimately shortens DS male lifespan, but lengthens DS female lifespan and helps them survive through the more stressful and resource-limited dry season (15, 16). While the behavioral ecology of these butterflies may explain seasonal variation in male courtship rates, the neural re-wiring that switches the male behavior is completely unknown.

In *B. anynana*, signaling of the hormone 20-hydroxyecdysone (20E) during early pupal development has been shown to regulate male courtship (17). In DS males, throughout pupal development, there are significantly lower levels of 20E titers circulating in the hemolymph than in WS males (17). The reduced courtship of DS males, however, can be switched to the WS active courting form by rearing pupae at higher temperatures during the first 50% of pupal development (15) or, alternatively, by keeping the pupae at low temperatures but injecting them with 20E at 30% of pupal development (17). These experiments suggest that this pupal stage is a critical period that determines male sexual behavior and that 20E may play an organizing role in the developing male brain of *B. anynana*. However, we currently have no direct evidence that 20E acts specifically on the brain to alter expression of genes that regulate male courtship behavior in adults. High levels of 20E could upregulate genes required for active courtship such as those described for *Drosophila* including *fruitless, doublesex* and *yellow* (9, 18, 19). Alternatively, 20E signaling may not lead to appreciable differences in the neural circuitry of dry and wet season male brains, but may instead influence other phenotypic traits that are also important in courtship behavior such as pheromone production (20) or the UV brightness of eyespot centers (16).

## Results

### Genes involved in melanin synthesis, dopamine metabolism and JH signaling are differentially expressed in pupal brains

To evaluate the direct organizational effects of 20E on the pupal brain of *B. anynana* males, and identify changes in gene expression levels that could impact the future sexual behavior of males, we compared the transcriptome of DS male brains injected with 20E at 30% pupal development (referred to as DS20E thereafter) with DS and WS male brains injected with a vehicle solution at the same developmental stage (DSV and WSV, respectively) (Fig. 1). RNA-seq extractions yielded a total of 2.11 x 10^8^ raw reads from 12 RNA-seq libraries. All read data was used to produce a transcriptome comprising of 1,403,420 total Trinity transcripts and 689,657 Trinity genes with an N50 length of 973 (full summary statistics are provided in Table S1 and Fig. S1). Mapping of the raw reads back to the transcriptome revealed that the overall alignment rate was 98.83-98.98%. Processing the transcriptome through CD-HIT (21) identified 1,045,896 unigenes in the transcriptome at 0.95 similarity. We identified 399 differentially expressed genes (DEGs) between DSV and WSV pupal brains, 302 were upregulated and 97 were downregulated in DSV. Comparing DS20E pupal brains to WSV, we identified 399 DEGs, 291 were upregulated and 108 were downregulated in DS20E. Comparing DS20E pupal brains to DSV we identified 151 DEGs, 79 were upregulated and 72 were downregulated in DS20E. Overall, the smaller number of differentially expressed genes observed between DSV and DS20E (151) compared to DSV and WSV (399) suggest that the DS20E brain transcriptome profile was more similar to DSV than WSV at 2 hours post-injection (Fig. S2 heat map). The full list of DEGs can be found in Table S2.

**Fig. 1.**
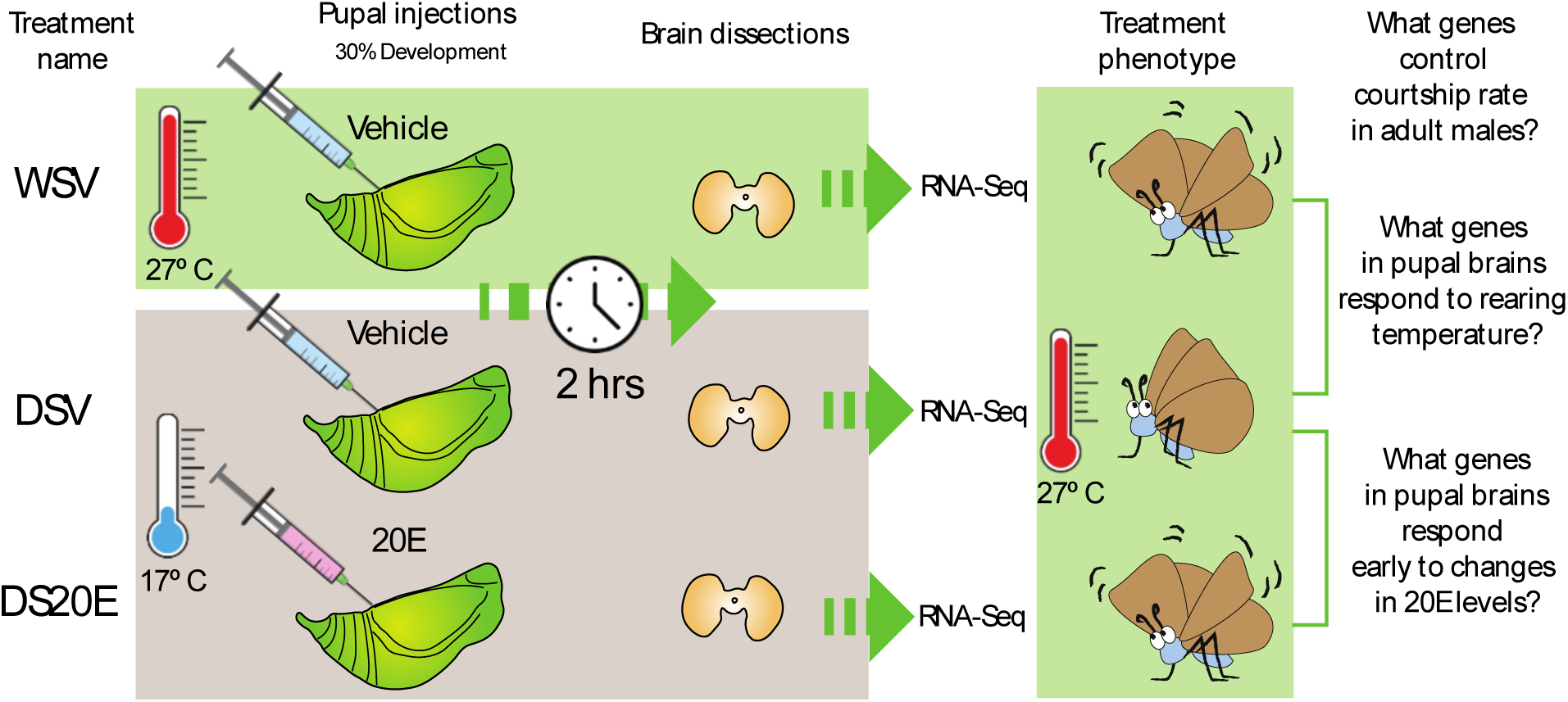
Overview of the experimental set-up for the pupal injections, brain dissections and RNA-seq for the three different treatment groups.

Genes known to be important in male courtship behavior such as *fruitless* and *doublesex* (22) were not differentially expressed at 30% of pupal brain development. However, the melanin pathway gene *yellow* was upregulated in both DSV and DS20E when compared to WSV. We also found that *dopamine N-acetyltransferase* (AANAT1/DAT1) was upregulated in DS20E (but not in DSV) when compared to WSV. Other potential genes of interest pertaining to courtship behavior included *Juvenile hormone* (JH), which was downregulated in DSV compared to WSV, and *Juvenile hormone epoxide hydrolase-like* (which hydrolyses JH) was upregulated in DSV and DS20E when compared to WSV. Genes involved in neural development included *neuropeptide CCHamide2, Neural Wiskott-Aldrich syndrome protein* and *lethal 2 essential for life l*(*2*)*efl*. These genes were downregulated in DSV compared to WSV with *l*(*2*)*efl* also downregulated in DS20E compared to WSV. Fig. 2 summarizes the log-fold change values and *p*-values of the top 10 annotated DEGs. See table S3 for a descriptive list of the DEGs associated with neural development.

**Fig. 2.**
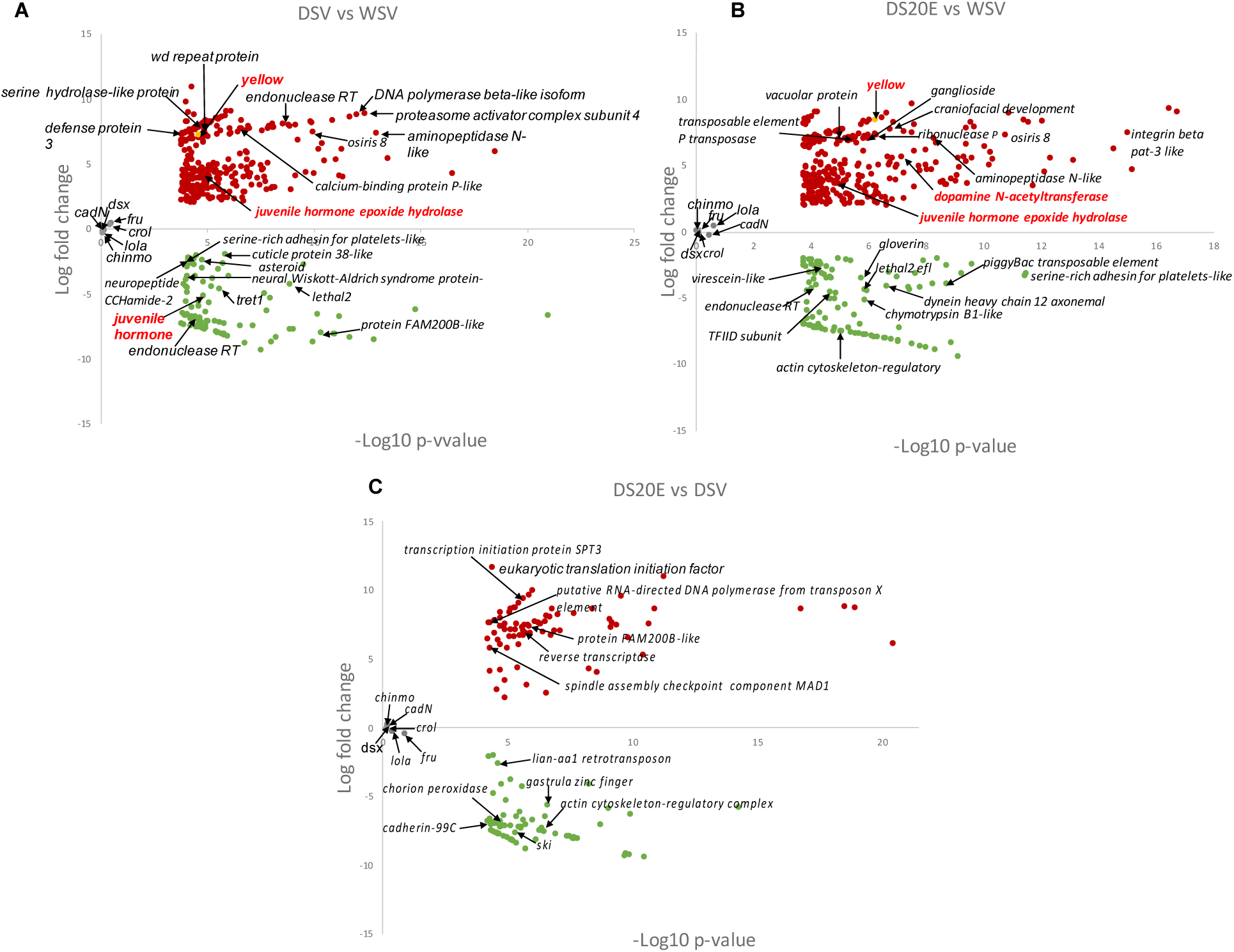
Volcano plots summarizing the log-fold change values and p-values of the differentially expressed genes. Annotations are shown for the top 10 differentially expressed genes which returned hits using the software Blast2Go (blastx to NCBI). Dopamine N-acetyltransferase (AANAT) and Juvenile epoxide hydrolase are also shown although they were not in the top 10. Upregulated genes are shown in red and downregulated genes are shown in green. **(A)** shows genes up- and downregulated in brains from the dry season vehicle treatment compared to the wet season vehicle treatment (DSV vs WSV). **(B)** shows genes up and downregulated in brains from the DS20E treatment (dry season pupa injected with 20E) compared to the WSV treatment (DS20E vs WSV). **(C)** shows genes up- and downregulated in brains from the DS20E treatment compared to DSV (DS20E vs DSV). Genes involved in male courtship behavior in *Drosophila* that were not differentially expressed in any comparison (grey).

### 20E downregulates the expression of yellow 4h after injections

From the transcriptomics analyses, we did not identify interesting candidate genes with a regulation pattern consistent with the high courtship of WSV and DS20E injected males, and the low courtship of DSV males. We thus decided to explore further the pupal brain expression of *yellow* because it was previously shown to affect courtship behavior in *Drosophila melanogaster* (18, 23) and was found to be upregulated in both DS treatments compared to WS. We hypothesized that 20E may affect *yellow* expression at a later stage post injection, impacting the courtship behavior in adults. To confirm the regulation of *yellow* expression by 20E in male pupal brains, we used qPCR to measure the relative expression of *yellow* in the brains of developing pupae at 3 different time points after injections of the hormone. Similar to the transcriptomics experiment, we injected 20E in DS male pupae at 30% development, and a vehicle solution in both DS and WS pupae at the same stage, and assessed the relative levels of *yellow* in dissected pupal brains at 2, 4, and 24 hours post injection.

Two hours post-injection, the levels of *yellow* were about 2.5 times higher in pupal brains of both DSV and DS20E compared to the level of expression in WSV pupal brains, (mirroring our RNA-seq results), although the expression levels were not significantly different (Fig. S3, ANOVA: F = 0.50, p = 0.62). However, at 4 hours post-injection, the expression of *yellow* increased significantly by about 8-fold in DSV compared to WSV pupal brains (Fig 3, ANOVA: F = 5.43, p =0.023; post-hoc analysis WSV-DSV: adj. p = 0.027), while the level of *yellow* expression in DS20E remained low, similar to those of WSV brains (ANOVA post-hoc analysis WSV-DS20E: adj. p = 0.79). Twenty-four hours post-injection, expression levels of *yellow* in the pupal brains of DSV and DS20E were respectively 2.8 and 3.7 times higher than in WSV (Fig. S3). Relative levels of *yellow* expression were significantly higher in DS20E than in WSV pupal brains (ANOVA: F = 4.12, p = 0.046; post-hoc analysis WSV versus DS20E: adj. p = 0.046). These results demonstrate that a single injection of 20E into DS pupae at 30% of development was sufficient to decrease *yellow* expression levels in DS20E to WSV levels at 4 hours post-injection. This single injection did not impact *yellow* levels at the earlier 2hr time period, nor keep *yellow* levels low at 24 hours post injection, suggesting that a short interval of time around 30% pupal development encompasses a hormone-sensitive window in which *yellow* is downregulated by 20E to WS levels.

**Fig. 3.**
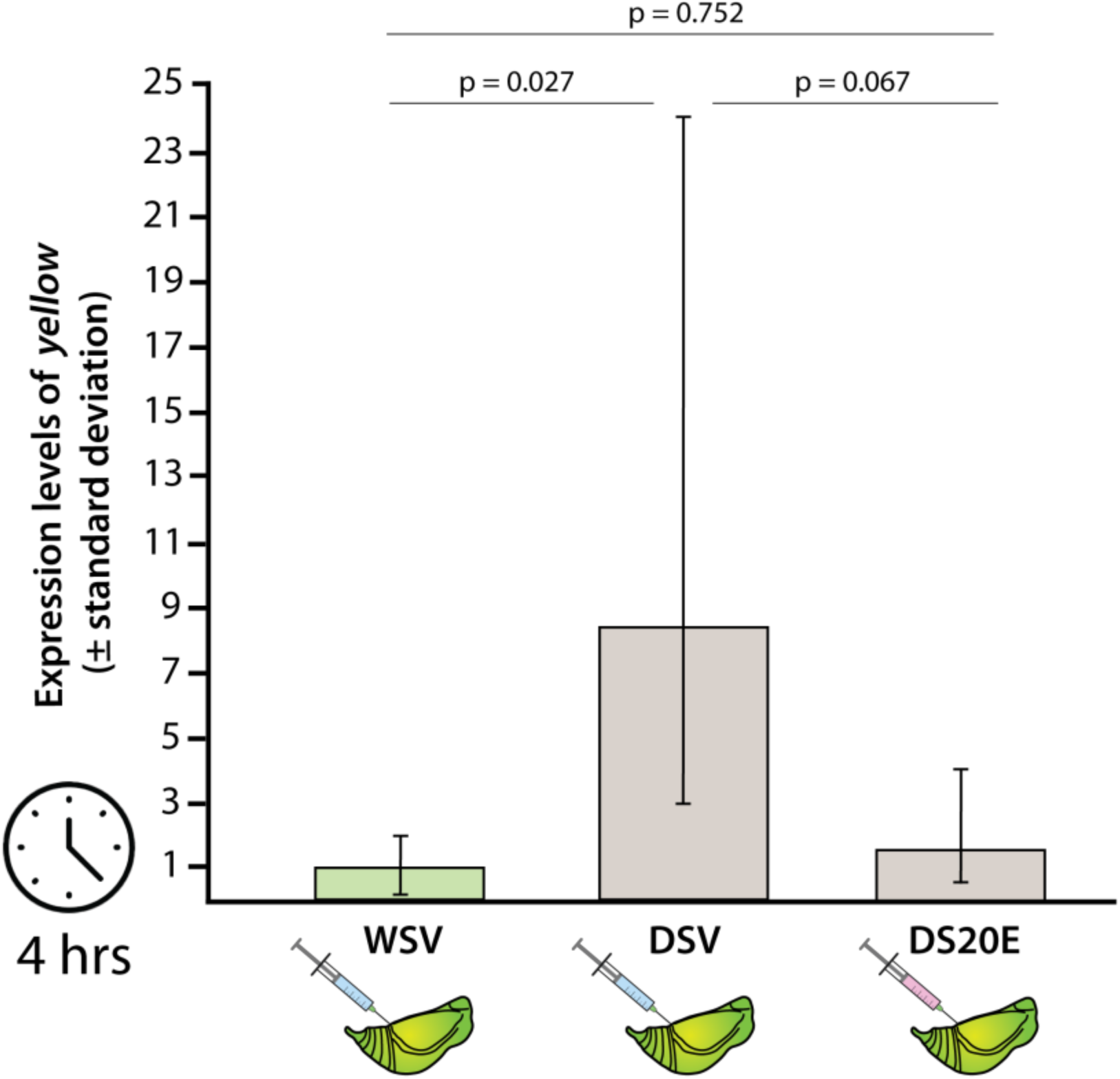
Four hours after injections, *yellow* is downregulated in brains of DS pupae injected with 20E compared to pupae injected with vehicle. Bars show fold change expression relative to WS pupae injected with vehicle solution. Indicated p values are the Turkey-adjusted p values from the post-hoc analysis.

### Yellow mutant males courted more frequently and for a longer duration than Wt males

We hypothesized that Yellow may be a repressor of male courtship as DS males exhibit lower courtship than WS males and have significantly higher expression of *yellow* during pupal brain development. To test this hypothesis, we generated a Yellow mutant homozygous line in *B. anynana* to investigate if loss of Yellow function leads to elevated levels of courtship in DS males alone or in both DS and WS males (Fig. S4). Using this Yellow knockout (KO) line, we compared the duration and frequency of the Yellow mutant and the wildtype (Wt) male courtship sequence, including copulation. Yellow mutant males courted for a longer duration (WS: t = 2.181, p = 0.0323; DS: t = 2.083, p = 0.0416; Fig. 4a) and more frequently (WS: z = 4.165, p < 0.0001; DS: z = 2.629, p = 0.00855; Fig. 4b) than Wt males regardless of seasonality. In addition, DS Yellow mutant males remained in copulation longer (t = 2.174, p = 0.039; Fig. 4c) than DS Wt males.

**Fig. 4.**
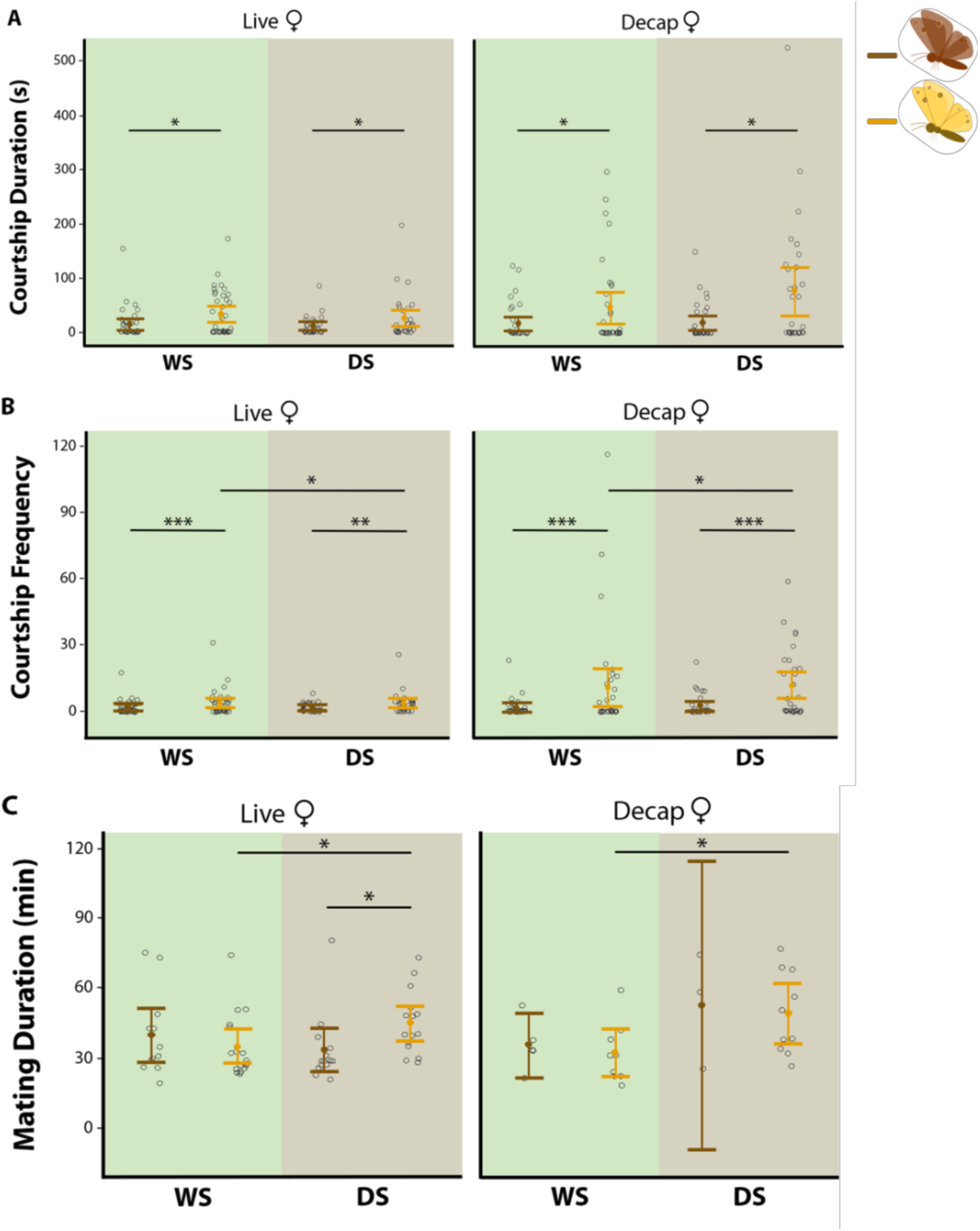
Courtship behavior of Wt and Yellow males, for wet (WS) and dry (DS) seasonal forms, in both live and decapitation assays. Yellow males courted at a higher duration and frequency than Wt males, for both WS and DS forms. DS Yellow males courted at a higher frequency and remained in copula for a longer period of time than WS Yellow males. **(A)** Courtship duration, **(B)** Courtship frequency and **(C)** Mating duration were quantified. Mating duration was quantified among mated males only. Vertical bars represent 95% confidence intervals. Open circles (°) are data points. Asterisks (*) indicate significant differences: * p ≤ 0.05, ** p ≤ 0.01, *** p ≤ 0.001. Outliers are not removed as they are true measurements. n(WS-Live-Wt) and n(WS-Live-Yellow) = 38, n(WS-Decap-Wt) and n(WS-Decap-Yellow) = 34, n(DS-Live-Wt), n(DS-Live-Yellow), n(DS-Decap-Wt) and n(DS-Decap-Yellow) = 30.

Because Yellow males have a lighter overall pigmentation, despite being identical in the brightness of a known sexual ornament, the eyespot’s ultraviolet reflecting white center on the ventral and dorsal sides of the forewing (Fig. S5) (24, 25), we repeated these male courtship observations using decapitated females. Decapitation prevents important visual cues detected by a female from impacting a male’s behavior, such as more intense courtship provoked by a female’s increased rejection behavior (26). Yellow mutant males still courted for a longer duration (WS: t = 2.083, p = 0.0416; DS: t = 2.269, p = 0.0266; Fig. 4a) and more frequently (WS: z = 10.59, p < 0.0001; DS: z = 8.246, p < 0.0001; Fig. 4b) than Wt males regardless of seasonality (see also Fig. S6). This result indicates that Yellow alters male courtship behavior independently of the female’s behavior towards those males.

### Yellow mutant males courted more frequently and copulated longer in their DS form than WS form

To test whether differences in Yellow expression levels were sufficient to explain courtship differences between the seasonal forms, we compared the amount of courtship between DS and WS Yellow mutant males. If Yellow, alone, was responsible for courtship differences between the forms then DS and WS Yellow mutant males should display similar levels of courtship. DS Yellow mutant males courted live females more frequently (z = 2.324, p = 0.0201; Fig. 4b) and copulated longer (t = 2.174, p = 0.039; Fig. 4c) than WS Yellow mutant males. Similar behavior was observed in males courting decapitated females, with DS Yellow mutant males courting more frequently (z = 2.454, p = 0.0141; Fig. 4b) and copulating longer (t = −2.34, p = 0.032; Fig. 4c) than their WS counterparts.

## Discussion

In insects, hormones are typically assumed to regulate sexual behavior by activating existing neural circuits that control processes such as sexual maturation, memory formation and pheromone communication (11, 27–29). Recently, it has been shown that hormones can also regulate sexual motivation by repressing activation of existing neural circuits (5). It remained unclear, however, whether exposure to different levels of hormones earlier in development could organize neural circuits that affect sexual behavior in adults. Here, we provide evidence that the ecdysteroid 20E plays an organizational role during pupal brain development in *B. anynana* by repressing expression of the *yellow* gene which leads to seasonal differences in male courtship behavior.

### Yellow is regulated by 20E during pupal brain development and functions a repressor of male courtship in B. anynana

We show that *yellow* is significantly upregulated in the pupal brains of DS male butterflies which court less than WS males. This increase in *yellow* expression appears to be in response to seasonal fluctuations in 20E, as injection of this hormone into DS males at 30% of pupal development was sufficient to suppress *yellow* expression to levels observed in WS pupal brains at 4 hours post-injection, as well as to rescue WS courtship levels in adults (17). These results suggest that this period encompasses a critical window during brain development which is sensitive to circulating levels of 20E. High levels of *yellow* in DS males suggested that *yellow* was a repressor of courtship. This was confirmed by knocking out *yellow* in *B. anynana* and observing male Yellow mutants exhibiting increased courtship frequency and duration compared to wildtype males of both seasonal forms. Given that Yellow WS mutants displayed more active courtship than wildtype WS males, this suggests that low levels of *yellow* expression are still required in wildtype WS males to reduce courtship and optimize energy expenditure, as increased wing fluttering observed in the Yellow KO line did not translate to increased mating success.

Comparing the behavior of Yellow mutant males between seasonal forms produced additional insights into the role of *yellow* in regulating male courtship plasticity. The complete loss of *yellow* led to DS males courting more than WS males. This result was surprising as it suggests that removal of Yellow inverts the relative amount of courtship performed by WS and DS males; It leads DS males to court more than WS males. This suggests that Yellow is required for inverting a biased level of courtship that would take place in these butterflies driven by temperature alone. Without the action of *yellow*, males reared at high temperature during development would court less than males reared at low temperatures. This indicates that the high levels of *yellow* expression in the brains of DS males is absolutely essential to produce the low levels of courtship in this seasonal form, and that other factors controlled by rearing temperature and by 20E are biasing adult courtship levels in the opposite direction to those observed in wildtype individuals. These factors can be explored in future.

Our findings are in striking contrast to those observed in *Drosophila* where *yellow* is required for normal male courtship behavior. Exactly how *yellow* expression influences male courtship behavior in *Drosophila* has been a topic of investigation that has yielded conflicting results. An early study suggested that *yellow* mutant males were less successful during courtship and displayed reduced wing vibrations (30). Further tests of these observations showed that mutations in *yellow* disrupted wing extension during the courtship ritual, preventing males from performing a courtship song which is required for male mating success (18, 31). However, recent work by Massey et al. argued that a lack of melanization in the sex combs of *yellow* mutants, rather than any impairment in neural circuitry affecting courtship song, was the trait that prevented males from successfully grasping females (19), an idea that was proposed earlier (32). It is possible that the fly laboratory stock might have evolved between the earlier and the later experiments, as the courtship observations repeated by Massey et al., produced different results from the original observations on the same stock (30). All research to date on *yellow* mutants in *Drosophila,* however, clearly demonstrate that *yellow* is absolutely required for successful male courtship.

### yellow may influence courtship behavior in B. anynana via the dopaminergic signaling pathway

Dopamine is an important catecholamine neurotransmitter which regulates a variety of behaviors including motor output, drive, arousal, pleasure and memory (33, 34). Dopaminergic signaling has been shown to regulate not only mating drive but also persistence and duration of mating in male *Drosophila* (35, 36). Melanin synthesis enzymes are expressed in the *Drosophila* brain and may be involved in the production of neuromelanin in dopaminergic neurons (37). Yellow is thought to function as a dopachrome conversion enzyme (DCE) in the melanin pathway converting L-Dopa to Dopa-melanin (38, 39). L-Dopa is also used as substrate for dopamine which is involved in both cuticle pigmentation and neurotransmission (36). Thus, variation in *yellow* expression could alter the availability of L-Dopa for dopamine synthesis, with higher expression of *yellow* in DS brains leading to reduced L-Dopa. Alternatively, Yellow may physically bind to dopamine, as demonstrated in a study of salivary proteins in sandflies (40). Thus, the increased expression of *yellow* in DS brains could lead to a reduction in dopamine availability, which may inhibit courtship behavior. In Fig. 5, we suggest a possible mechanism of Yellow involvement in the pathway converting tyrosine to L-Dopa in dopaminergic neurons.

**Fig. 5.**
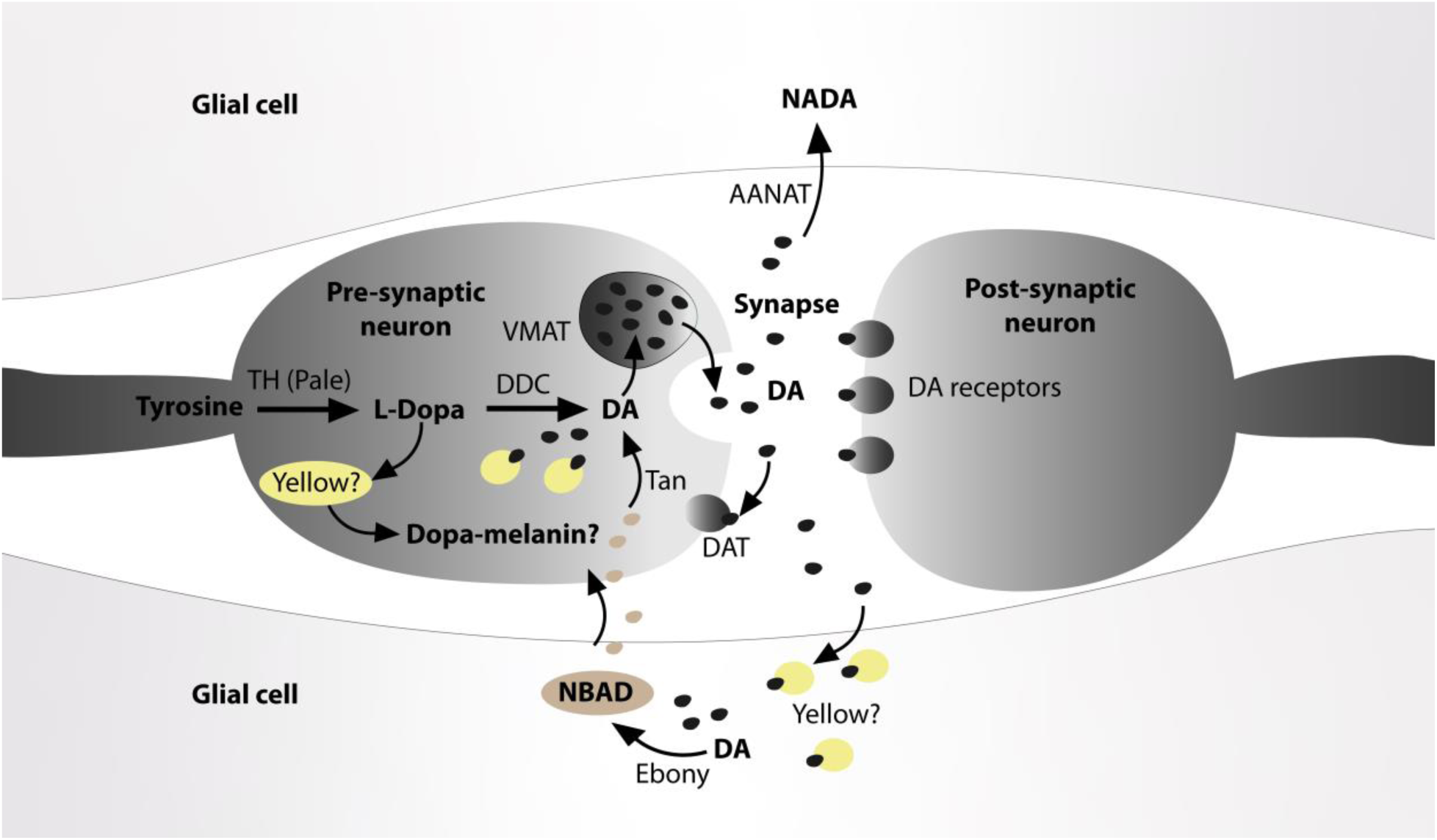
Schematic model of a melanized dopaminergic neuron adapted from Yamamoto and Seto (36). In this model tyrosine is converted to L-Dopa which is converted to dopamine (DA) by Dopa decarboxylase (DDC). We hypothesize that *yellow* may also be expressed in dopaminergic neurons to convert L-Dopa to Dopa-melanin. Upregulation of *yellow* in DS pupal brains could decrease availability of L-Dopa to produce dopamine which is packaged into vesicles (VMAT) for release into the synapse. Another possibility is that Yellow binds to dopamine, reducing its availability either for direct synthesis of dopamine by DDC or indirectly (if it is also expressed in glial cells) by limiting the availability of dopamine conversion to NBAD by Ebony, which is converted back to dopamine by Tan. An excess of DA is converted to NADA by AANAT or is taken up by DAT back into the pre-synaptic neuron.

Currently we have no direct evidence that dopamine levels differ between DS and WS *Bicyclus* brains. However, the expression profile of a few genes in our transcriptome analyses may provide some indirect evidence. We found that dopamine N-acetyltransferase, AANAT/DAT1 was upregulated in pupal brains of DS20E (but not in DSV) as compared to WSV brains. This may indicate a transient response to the 20E injection. The function of AANAT is to metabolize and inactivate secreted dopamine in the synapse shortly after release (33, 37). In young female *Drosophila virilis*, higher titers of 20E leads to an increase in dopamine, although this appears to be associated with reduced activity of AANAT (41). However, in retinal cells of fish, AANAT activity is positively correlated with dopamine levels (42). An increase in dopamine induced by 20E would provide a mechanistic explanation as to why DS male pupa injected with 20E display active WS courtship behavior. Functional experiments measuring or manipulating dopamine levels, however, are required to test the hypothesis that dopamine levels are higher in WS male pupal brains.

We also observed changes in Juvenile hormone (JH) signaling, which is known to interact with dopamine to affect sexual maturity and courtship behavior in *Drosophila,* likely through changes in neural development (14, 43). In DS pupal brains, JH was downregulated and Juvenile hormone epoxide hydrolase (JHE), which degrades JH (44), was upregulated. These findings could indicate that dopamine levels are low in DS male pupal brains as dopamine increases JH titers in young female *D. virilis* by inhibiting its degradation (45). JH is also associated with increased dopamine levels in male honeybees (46). Although we also see an upregulation of JHE but no longer a downregulation of JH in DS20E, this may reflect a response to changing levels of dopamine. Interactions between 20E, JH, dopamine and AANAT in *Drosophila* represent a complex pathway as depicted in (41, 47) thus, we must interpret our findings with caution. However, given that this is an important pathway for regulating courtship behavior in *Drosophila*, the differential expression of these genes in our transcriptome analyses suggest their possible involvement in regulating courtship behavior in *B. anynana*.

### Drosophila courtship genes do not impact Bicyclus courtship behavior at this pupal stage

A number of genes that are known to be important for male courtship in *Drosophila* (*fruitless, dsx crol, lola, cadN* and *chinmo*) (22), were surprisingly not differentially expressed between any of our treatment groups and all showed very low levels of expression (Fig. 2). Thus, it appears that in *B. anynana* these genes do not influence neural re-wiring occurring at this particular stage of brain development. However, we cannot exclude the possibility that these genes are upregulated at a later time point after the 20E injection. We did, however, identify some other differentially expressed *genes* including *Neural Wiskott-Aldrich syndrome protein* (N-WASP), *CCHamide2* and *lethal 2 essential for life (l*(*2*)*efl)* that may be important in neural development and behavior and could serve as interesting candidates for future studies (see Table S3).

## Conclusions

Here, we show that male DS butterflies court less than WS butterflies due to temperature-induced changes in levels of 20E, which alter the expression of the *yellow* gene during pupal brain development leading to differences in adult courtship behavior. In *B. anynana*, *yellow* functions as a repressor of male courtship and is a downstream target of 20E. Our results suggest potential interactions of 20E on JH and dopamine signaling, a circuit that is well described in *Drosophila*. Future studies examining dopamine levels between the seasonal forms and the individual role of 20E and JH on dopaminergic signaling would help clarify mechanistically why the *yellow* gene functions as a repressor of male courtship in these butterflies. We propose an organizational role for 20E, suggesting convergence in hormone regulation of sexual behavior in insects and vertebrates. For animals living in seasonal environments, selection may favor adaptations that use external cues to optimize behavior, such as employing environmentally induced hormones like 20E to organize neural circuits during critical windows of brain development.

## Acknowledgements

We would like to acknowledge Dong Qiang Cheng, for helping EJT with the hardware issues on lab server; Jan Gruber, for advising EJT with the Linux operating system and also Firefly Farms in Singapore for supplying corn for larval rearing.

## Methods

### Transcriptome assembly and analyses

#### Sample collection and Illumina(R) RNA-seq experiments

To mimic dry and wet season conditions, caterpillars of *B. anynana* were reared under WS and DS temperatures in climate-controlled rooms at 27°C and 17°C, respectively, at 80% humidity, and a 12:12 hr light:dark photoperiod. Caterpillars were fed corn plants *ad libitum* until pupation. Pupae were staged, such that the percent of pupal development was known for all individuals. At 30% of pupal development (day 2 in the WS butterflies and day 6 in the DS butterflies) DS pupae were injected with either 3 μl of 2000 pg/μl (6000 pg total) (10% 20E in EtOH + 90% saline) of 20E (Sigma-Aldrich®) or with 3 μl of vehicle (10% EtOH and 90% saline) and WS pupae were injected with 3 μl of vehicle in the lateral posterior region of the fifth abdominal segment. The injections were done at 1200 h and the brains were dissected two hours later, at 1400 h. We chose to inject the animals two hours before collection in order to give the 20E time to circulate through the open circulatory system of the insect to reach the brain and to affect gene expression in this tissue. We chose this time point after injection to collect the samples because previous studies have demonstrated that genes, which respond early to 20E signaling are expressed about 2 hours after exposure to 20E (48, 49).

Samples from each treatment were collected on each collection day in order to reduce the confounding effects of day of collection on gene expression. Each sample consisted of three biological replicates of wet season pupal brains following treatment with vehicle only; four biological replicates of dry season pupae treated with vehicle only; and five biological replicates of dry season pupae treated with 20E. Each biological replicate was made up of mRNA pooled from the brains of five different individual male pupae. Brains were pooled to account for the genetic variation in the colony. The pupal brains were dissected in a solution of ice cold 1X PBS. After each dissection, the brain was immediately immersed in a 1.5 eppendorf tube containing 500 μl of TRIzol® reagent (Life Technologies®) and 3 RNase-free beads (#SSB14B 1.4mm Stainless Blend NEXT>>>ADVANCE®).

Once all five brains from a particular treatment group were placed in the TRIzol reagent, the brains were immediately homogenized using a Bullet Blender (NEXT>>>ADVANCE®) for three minutes and total RNA was extracted using the trizol-chloroform protocol. DNA was removed using gDNA Eliminator Mini Spin Columns from the RNeasy Plus Micro Kit (Qiagen®) and following the kit instructions. The quality of the extracted RNA was checked using a ND1000 spectrophotometer (NanoDrop® Technologies) and stored at −80°C. The samples were submitted to the W.M. Keck Biotechnology Resource Laboratory for Illumina® RNA-Seq. The RNA samples consisted of 4 μg of total RNA in 20 ml of water, and was run on a separate lane of a flow cell on a HiSeq2000. The Keck Biotechnology Resource processed the samples following standard Illumina® RNA-Seq protocol.

#### Transcriptome assembly

We assembled the transcriptome from a total of 12 RNA-Seq libraries. Raw reads of the RNA-seq libraries were uploaded to the SRA database with the SRA accession number PRJNA544388. Prior to performing the transcriptome assembly, we performed quality trimming of the input raw reads using Trimmomatic using the default options (50). We assembled a de novo transcriptome using Trinity 2.4.0 (51) and Bowtie2, following the protocol by Haas et al. (52). The transcriptome was then uploaded to the Transcriptome Shotgun Assembly (TSA) Database, following the TSA guidelines. During this process, transcripts were screened for vector contaminations and any vector and linker sequences were removed. In addition, transcripts smaller than 200 bp were screened and removed from the assembly. This Transcriptome Shotgun Assembly project has been deposited at DDBJ/EMBL/GenBank under the accession GHRJ00000000. The version described in this paper is the first version, GHRJ01000000.

To characterize the quality of our transcriptome assembly, we used scripts in the Trinity toolkit. First, we computed assembly statistics, which is the contig N50 value based on the set of transcripts representing 90% of the expression data, using the TrinityStats.pl script. These assembly statistics for *B. anynana* brain transcriptome are reported in Table S1. Next, we computed the N50 statistics of the top most highly expressed transcripts that represent x% of the total normalized expression data, using the contig_ExN50_statistic.pl script. The N50 statistics are presented in Figure S1. To assess the proportion of raw reads mapped to the transcriptome assembly, we used Bowtie2. We then extracted transcripts that are most differentially expressed and clustered these transcripts according to their patterns of differential expression using the analyze_diff_expr.pl script from the Trinity toolkit. The clustering analysis indicated that the biological replicates from the same treatment clustered together, as shown in Figure S2.

Next, we obtained unigenes for the transcriptome using CD-Hit version 4.6 (21) with similarity set to 0.95. CD-Hit clustered all sequences with similarity ≥ 95%, retaining only the longest transcript, thus splice variants/isoforms were removed and redundancy was reduced. The transcriptome assembly with unigenes was then used for differential gene expression, described in the next section.

#### Differential gene expression

To estimate transcript abundance, we used RSEM 1.3.2 (53), a software which uses Bowtie2 (54) to align the transcripts to the transcriptome assembly, thus quantifying gene and isoform abundances. The RSEM output reported normalized expression metrics as fragments per kilobase transcript length per million fragments mapped (FPKM) and transcripts per million transcripts (TPM). Next, we used edgeR 3.28.1 (55), to examine differential expression of genes across the three treatments (DSV vs WSV, DS20E vs DSV and DS20E vs WSV). EdgeR normalizes RNA composition by finding a set of scaling factors for the library sizes that minimize the log-fold change ratios (logFC) between the samples for most genes. We used the default method in edgeR for computing these scale factors, which is a trimmed mean of Mvalues (TMM) between each pair of samples. Genes with a False Discovery Rate (FDR) of <0.05 and logFC of ≥2 were defined as differentially expressed genes (DEGs).

In order to understand the DEGs induced by the hormone treatment, we further annotated the DEGs with Blast2GO 5.2.5. We used the public NCBI Blast service (QBlast) to blast our sequences against the non-redundant protein database using the blastx-fast program. Matched transcripts were filtered using a cut-off E-value of 1 x 10^−3^; otherwise the default settings for Blast2GO were used at each step. To annotate the remaining transcriptome, we performed a local blastx of the assembled contigs against the *Bicyclus anynana* v1.2 draft genome (56). The annotated transcriptome has been deposited on Dryad and will be made available for publication.

#### qPCR sample collection and experiments

Sample collection was similar to the one described above for the RNA seq experiment. We measured the total development time of Wt pupae, and at 30% development (2.5 and 6.5 days for WS and DS pupae respectively in these rearing conditions), we injected pupae with 20E or vehicle solutions using the same protocol as described above. Pupal brains were dissected 2 hours, 4 hours and 24 hours after injections in ice cold 1X PBS, placed immediately into RNALater (Qiagen, GmbH, Hilden, Germany) and stored at −20°C until RNA extraction. We used 5 biological replicates per treatment, each made of 5 pooled brains (for the 2 hours dissections) or 2 pooled brains (for the 4- and 24-hour dissections).

Total RNA extraction, including the elimination of genomic DNA, was done using the Qiagen RNeasy Plus Mini Kit (Hilden, Germany) following the manufacturer’s instructions. Complementary DNA was synthesized using the RevertAid RT Reverse Transcription Kit (Themoscientific). 10 ng of cDNA were used for qPCR with the KAPA SYBR FAST qPCR Kit (KK4604, KAPA Biosystems, Wilmington, MA, USA) and the experiment run on the Biorad CFX96 system using the TqPCR protocol described in Zhang et al. (57). Primer efficiencies were calculated using 0.5, 5 and 50 ng of cDNA from Wt tissue (with 3 technical replicates). The primers are described in Table S4.

We calculated the relative transcript levels using the common based method (58). The Ct values were normalized to the reference gene *EF1α* and to the average Ct of the WS reference samples. ΔCt values from each treatment were compared using a one-way analysis of variance (ANOVA) followed by a post-hoc analysis providing p values adjusted with the Tukey method. Statistical analyses were performed in R v.4.0.0 (59) implemented in RStudio v.1.2.5042 (60), using Rmisc, car and emmeans packages (61–63).

#### Generation of CRISPR-attP Yellow knock out line

To establish a Yellow mutant line we inserted an attP sequence into exon 4 of the *yellow* gene to disrupt its overall sequence (Fig S7). We used a knock-in method through homology directed repair (HDR) using a single-stranded DNA (ssDNA) as a template. The ssDNA construct was made following methods described in (64). The ssDNA contains 66 bp and 60 bp of homologous sequence around the target region on each side of the attP sequence motif. We injected 500 ng/μl of a sgRNA targeting the *yellow* gene, 500 ng/μl Cas9 mRNA, and 160 ng/μl ssDNA into fertilized 2-3 hr old embryos. Out of 254 injected embryos, 87 larvae hatched, resulting in 14 adults (6 males and 8 females). We then crossed 3 G^0^ mosaic butterflies (showing some yellow patches of coloration on the wings) with 3 Wt to obtain the G^1^ generation. To identify which cage contained transgenic butterflies with the attP insertion, we pooled 30 embryos from each cage, extracted genomic DNA, and performed PCR. One out of 3 cages showed a positive band. In cages that were identified as contained transgenic individuals we performed further genotyping of individual larvae using haemolymph PCR (Fig. S7). We isolated 5 transgenics out of 102 G^1^ genotyped animals. We confirmed that the PCR amplicon flanking the gRNA target site was the expected sequence although it contained 2 substitutions outside the attP sequence (Fig. S7). We crossed these 5 positive G^1^ butterflies with a Yellow phenotype with Wt counterparts, and then identified heterozygous G^2^ mutant offspring by haemolymph PCR since *B. anynana yellow* gene is likely a dominant gene regarding its effect on body color. We further crossed heterozygous G^2^ butterflies with each other, and obtained a homozygous G^3^ generation using, again, genotyping via haemolymph PCR.

### Behavioral Assays

#### Animal husbandry

Larvae from both the Yellow CRISPR-attP line and Wt were fed with young maize plants (*Zea mays*) and adults with mashed bananas *ad libitum*. Larvae of both lines were reared in WS and DS conditions as described above. Prior to eclosion (Day 0), pupae were separated according to their sex to ensure virginity. Adults that emerged on the same day were then transferred to other cages and dated accordingly.

#### Behavioral experiments

Behavioral assays were conducted in cylindrical hanging cages (30cm x 40cm) under one full spectrum light tube (Plantmax) and one UV light bulb (Arcadia Marine Blue), at 23°C, from 17:00 to 18:00. This specific time of observation was chosen because *B. anynana* exhibits crepuscular courtship (17). Visual barriers were placed between cages to prevent mate-copying (65). Within each sex, butterflies used for each assay were of the same age. All butterflies used in the assays ranged from four to eight days old. Two experiments were performed, one with live females and the other with decapitated females. The treatments were i) two Wt males x two Wt females, and ii) two Yellow mutant males x two Wt females (Fig. S4). One of the two males/ females in an assay was dotted with a black marker at both of its ventral hindwings to allow for sex-specific scoring of behavior. The multiple elements of courtship, as documented in Nieberding et al. (66), were scored in the assays: 1) localization (flying to other butterfly), 2) rapid flickering of wings, 3) thrusting (touching female’s wings with head), and 4) attempting (curling of the abdomen) (Fig. S4). Orientation (orienting body to female’s posterior) was not recorded since it was difficult to score or interpret their intent (courtship or coincidence) with that behavior. Latency to mate (time taken from the start of assay to the first mating) and mating duration were recorded as well. Behavioral assays lasted one hour, and quantification of an individual male behavior stopped once the first mating had occurred (i.e. even if one has mated, the other male’s behavior is still quantified until its own mating or one hour has lapsed).

For decapitation experiments, only females were decapitated to characterize male sexual behavior in the absence of female response (26). The same behavioral scoring as described above was used for this experiment. Decapitated females were first anesthetized in a −20°C freezer for 20 minutes and their heads were removed. Females were pinned through their thorax into opposite sides of the cage as illustrated in Fig. S4c and d. Based on personal observations, males tend to begin courtship when females start moving or signal readiness. Without movement, which was observed in some of the decapitation assays, males do not attempt to court at all, regardless of treatment type. Therefore, to overcome this potential issue, the pins (and therefore the thorax) of decapitated females were moved gently to emulate movement after 30 minutes had passed since the start of the assay. This specific timepoint was used as the average mating latency for live experiments was approximately 30 minutes. Thus, this reduces bias as much as possible, while still managing to test the effect of the *yellow* gene in male courtship behavior.

#### Statistical analysis

The data was evaluated for equality of variances and normality using the Levene’s test and Shapiro-Wilk test, respectively. Total duration and frequency of courtship were calculated by adding up the duration/ frequency of all the courtship elements displayed (localising + flicker + thrust + attempt) during the observation period. A Generalised Linear Model (GLM) with a Tweedie distribution (Gamma family; tweedie package (67)) was done to test the impact of the treatment type (Yellow/ WT) and season (WS/ DS) on the total duration of courtship. A Tweedie distribution (Gamma) was used due to the high number of zeroes and the skewness of data. The impact of the treatment type (Yellow/ WT) and season (WS/ DS) on the frequency of each courtship element was compared using a Zero-Inflated Poisson Model (pscl package (68)). Both mating latency and mating duration of the mated pairs were compared using independent t-tests. Chi-square tests were carried out to identify any associations between treatment type and mating success. Statistical tests and figures were done with IBM SPSS Statistics 25 and R-4.0.2 (59). The spectral data of eyespots was visualized using the pavo package (Fig. S5 (69)). Spectral analysis was done through calculating area under curve (AUC) for each eyespot replicate and the AUC analyzed using an ANOVA with post-hoc Tukey test in R.

## Supplementary data and information

### Supplementary tables

**Table S1.** Assembly statistics for *B. anynana* brain transcriptome

| Type | Assembly statistics |
| --- | --- |
| Total Trinity genes | 689,657 |
| Total Trinity transcripts | 1,403,420 |
| Percent GC | 36.50 |
| Median contig length (bp) | 456 |
| Average contig length (bp) | 703.17 |
| N50 (bp) | 973 |
| Total assembled bases | 986,848,704 |

**Table S2 (excel file).** Differentially expressed genes of male pupal brains comparisons between treatments (Dry season with vehicle - DSV, Wet season with vehicle - WSV and Dry season injected with 20E - DS20E. logFC represents the log-foldchange in the gene expression; logCPM represents the log counts per million; FDR represents the false discovery rate, where values of less than 0.001 are simply represented as <0.001. **See excel file** sheet 1 for DSV compared to WSV, sheet 2 for DS20E compared to DSV, and sheet 3 for DS20E compared to WSV. Green color indicates down-regulated genes and red indicates upregulated genes.

**Table S3.**
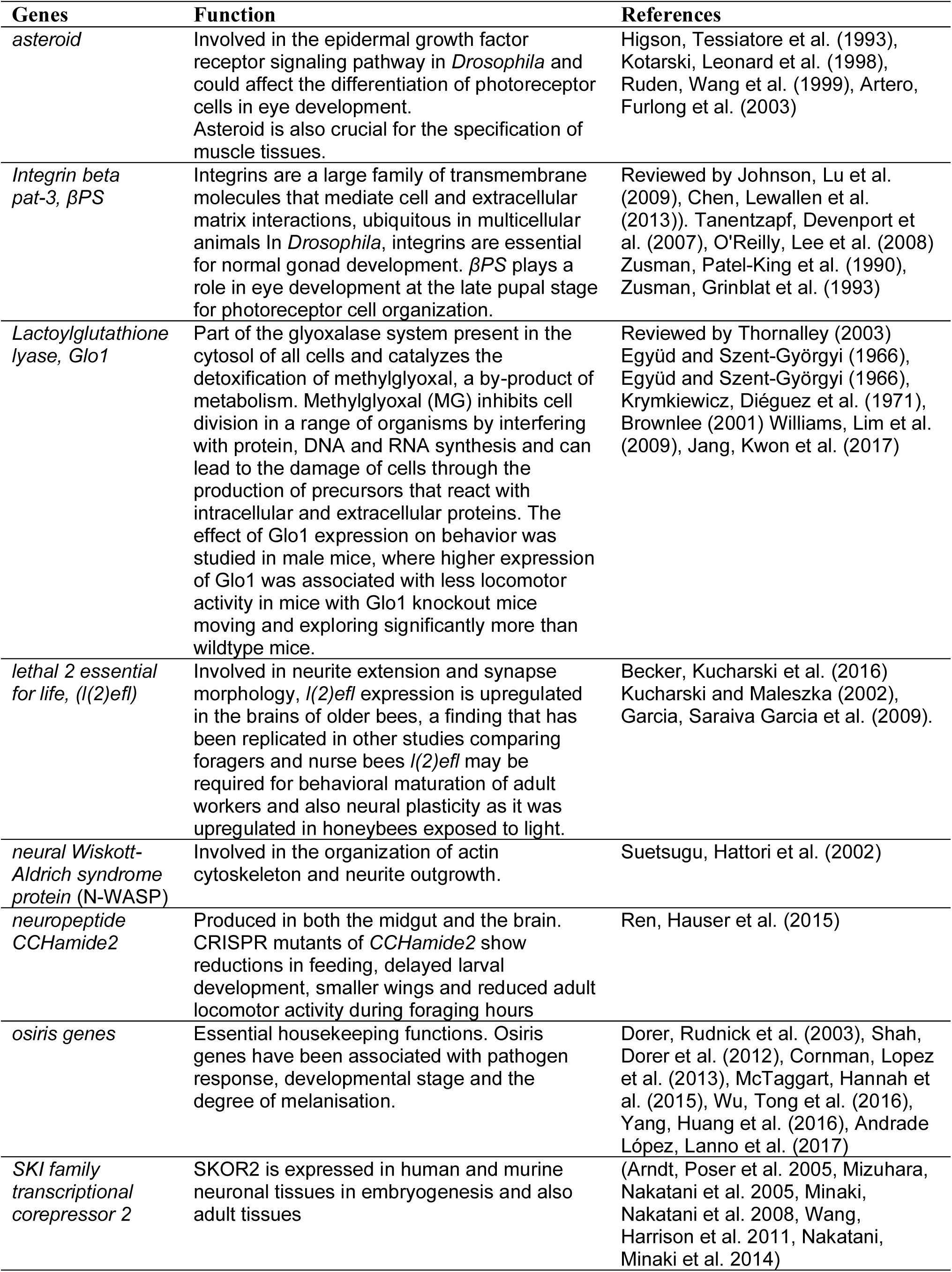
Differentially expressed genes associated with neural development.

| **Genes** | **Function** | **References** |
| --- | --- | --- |
| *asteroid* | Involved in the epidermal growth factor receptor signaling pathway in *Drosophila* and could affect the differentiation of photoreceptor cells in eye development. Asteroid is also crucial for the specification of muscle tissues. | Higson, Tessiatore et al. (1993), Kotarski, Leonard et al. (1998), Ruden, Wang et al. (1999), Artero, Furlong et al. (2003) |
| *Integrin beta pat-3, βPS* | Integrins are a large family of transmembrane molecules that mediate cell and extracellular matrix interactions, ubiquitous in multicellular animals. In *Drosophila*, integrins are essential for normal gonad development. βPS plays a role in eye development at the late pupal stage for photoreceptor cell organization. | Reviewed by Johnson, Lu et al. (2009), Chen, Lewallen et al. (2013)). Tanentzapf, Devenport et al. (2007), O'Reilly, Lee et al. (2008) Zusman, Patel-King et al. (1990), Zusman, Grinblat et al. (1993) |
| *Lactoylglutathione lyase, Glo1* | Part of the glyoxalase system present in the cytosol of all cells and catalyzes the detoxification of methylglyoxal, a by-product of metabolism. Methylglyoxal (MG) inhibits cell division in a range of organisms by interfering with protein, DNA and RNA synthesis and can lead to the damage of cells through the production of precursors that react with intracellular and extracellular proteins. The effect of Glo1 expression on behavior was studied in male mice, where higher expression of Glo1 was associated with less locomotor activity in mice with Glo1 knockout mice moving and exploring significantly more than wildtype mice. | Reviewed by Thornalley (2003) Együd and Szent-Györgyi (1966), Együd and Szent-Györgyi (1966), Krymkiewicz, Diéguez et al. (1971), Brownlee (2001) Williams, Lim et al. (2009), Jang, Kwon et al. (2017) |
| *lethal 2 essential for life, l(2)efl* | Involved in neurite extension and synapse morphology, *l(2)efl* expression is upregulated in the brains of older bees, a finding that has been replicated in other studies comparing foragers and nurse bees *l(2)efl* may be required for behavioral maturation of adult workers and also neural plasticity as it was upregulated in honeybees exposed to light. | Becker, Kucharski et al. (2016) Kucharski and Maleszka (2002), Garcia, Saraiva Garcia et al. (2009). |
| *neural Wiskott-Aldrich syndrome protein (N-WASP)* | Involved in the organization of actin cytoskeleton and neurite outgrowth. | Suetsugu, Hattori et al. (2002) |
| *neuropeptide CCHamide2* | Produced in both the midgut and the brain. CRISPR mutants of *CCHamide2* show reductions in feeding, delayed larval development, smaller wings and reduced adult locomotor activity during foraging hours | Ren, Hauser et al. (2015) |
| *osiris genes* | Essential housekeeping functions. Osiris genes have been associated with pathogen response, developmental stage and the degree of melanisation. | Dorer, Rudnick et al. (2003), Shah, Dorer et al. (2012), Cornman, Lopez et al. (2013), McTaggart, Hannah et al. (2015), Wu, Tong et al. (2016), Yang, Huang et al. (2016), Andrade López, Lanno et al. (2017) |
| *SKI family transcriptional corepressor 2* | SKOR2 is expressed in human and murine neuronal tissues in embryogenesis and also adult tissues | (Arndt, Poser et al. 2005, Mizuhara, Nakatani et al. 2005, Minaki, Nakatani et al. 2008, Wang, Harrison et al. 2011, Nakatani, Minaki et al. 2014) |

**Table S4.**
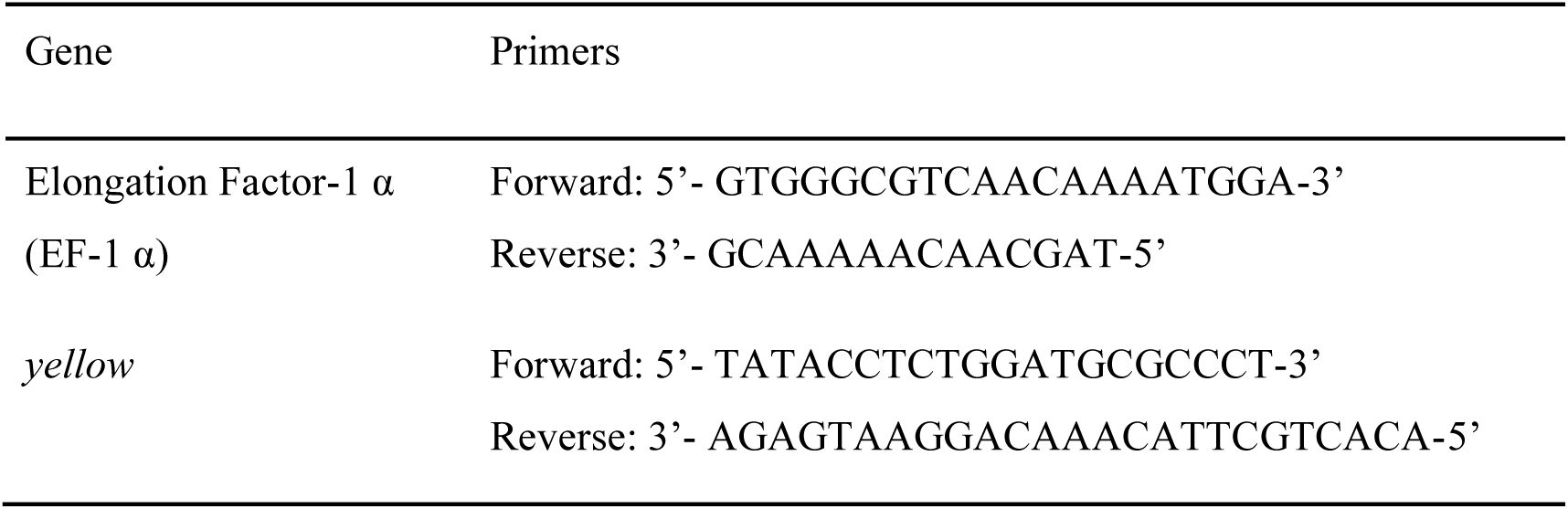
Primers used in the qPCR experiment

### Supplementary Figures

**Fig. S1.**
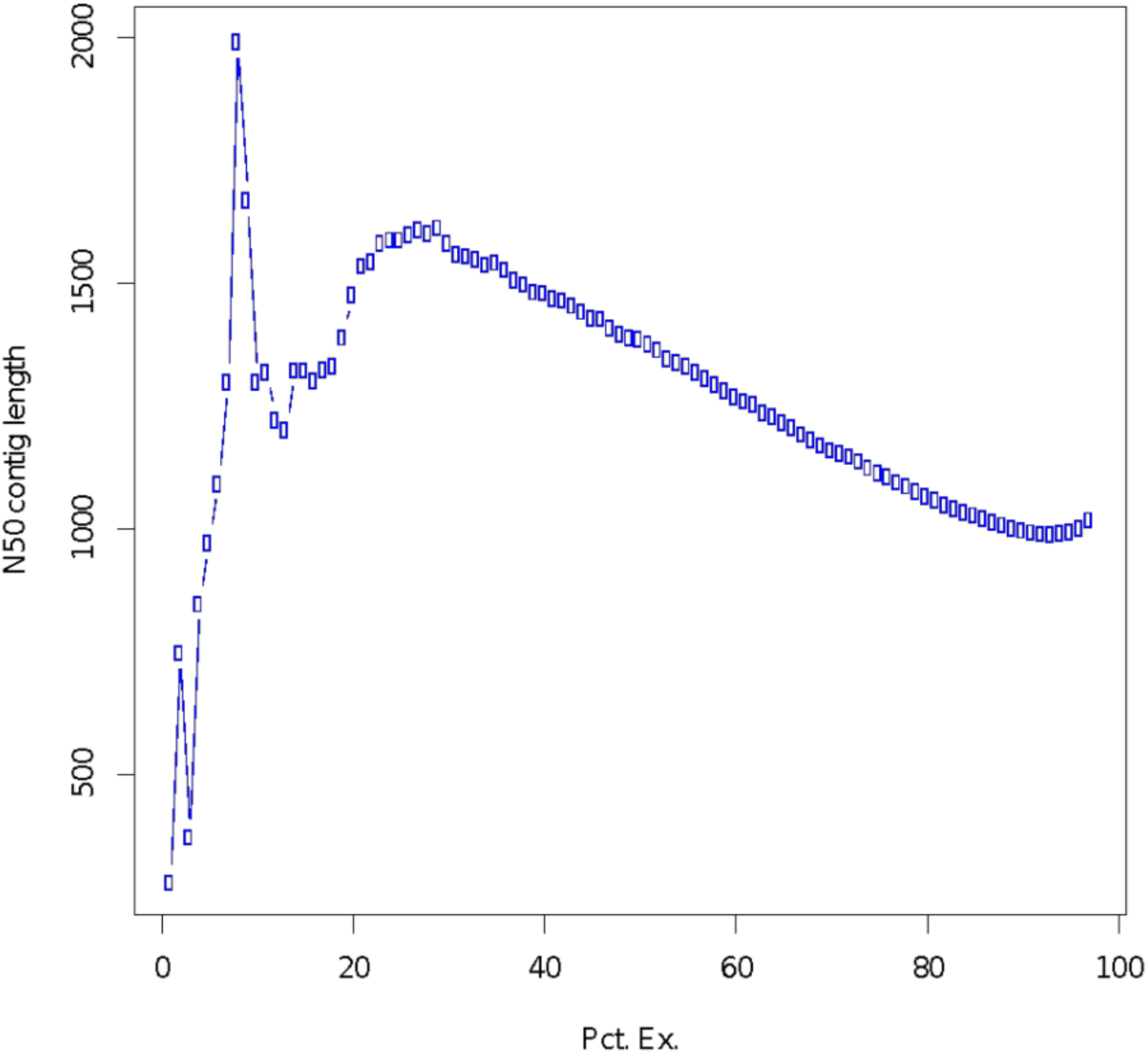
Summary of N50 contig lengths against ExN50 values

**Fig. S2.**
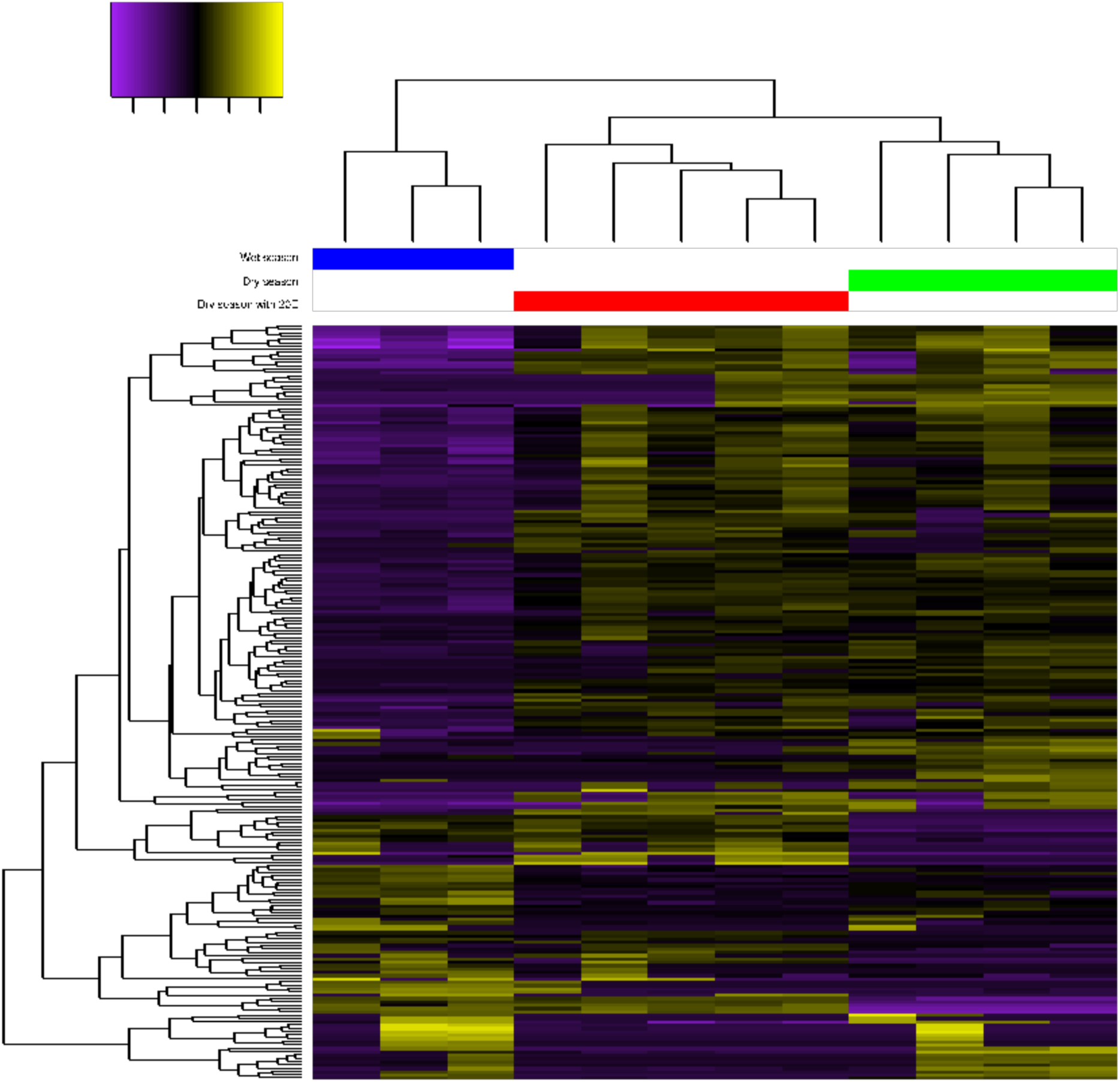
Clustering analysis of transcripts across 12 libraries. Colored bars at the base of the dendrogram indicate libraries of the three treatments – blue bar: wet season (S1, S12, S17); red bar: dry season with 20E (S2, S3, S6, S8, S15) and green bar: dry season (S4, S5, S7, S9). Dendrogram on the left represents the clustering of transcript expression based on the frequency of their expression together. The color of transcripts indicates their relative expression. The color key indicates the range of colors representing the fold change difference in expression.

**Fig. S3.**
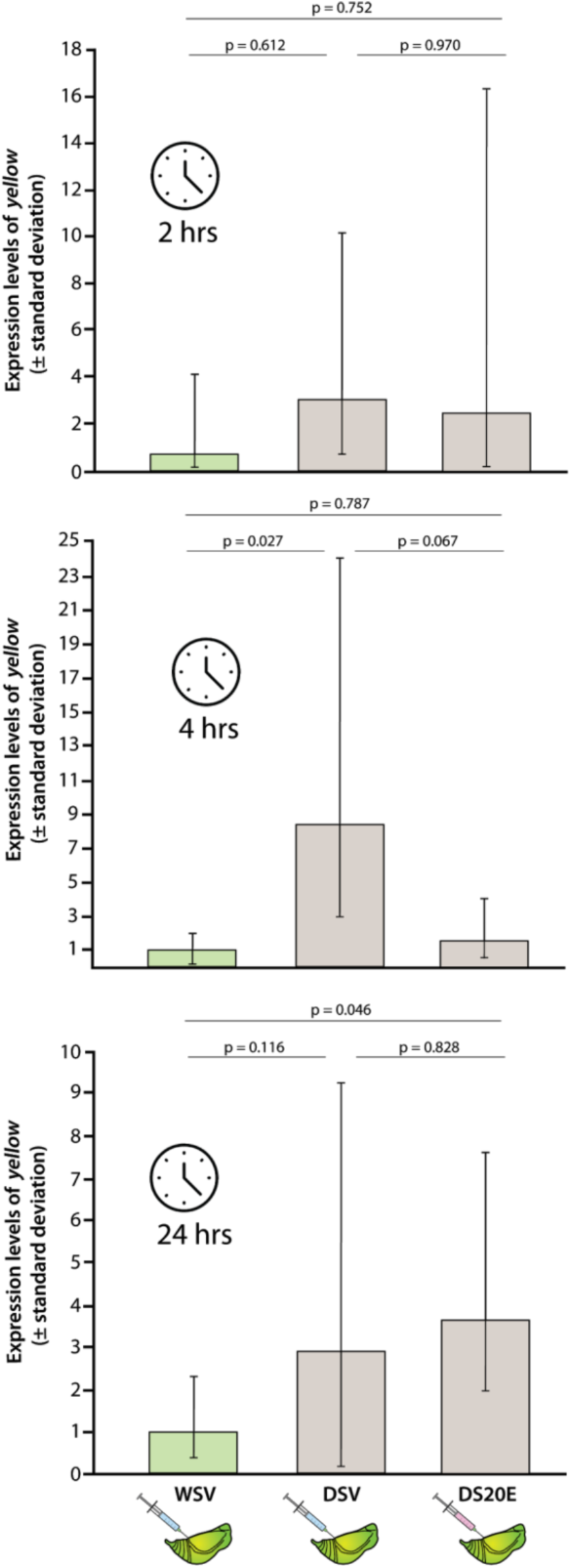
Expression level of yellow in brains of vehicle- and 20E-injected DS pupae dissected 2 hours, 4 hours and 24 hours after injections. Levels of expression are calculated relative to the baseline groups, all WS pupae injected with vehicle solution (at the value of 1, from 5 biological replicates at 2h and 4 replicates at 4 and 24 h) and dissected at the same time as DS individuals (5 biological replicates in all DS groups). Indicated p are the Tukey adjusted p values from the post-hoc analysis).

**Fig. S4.**
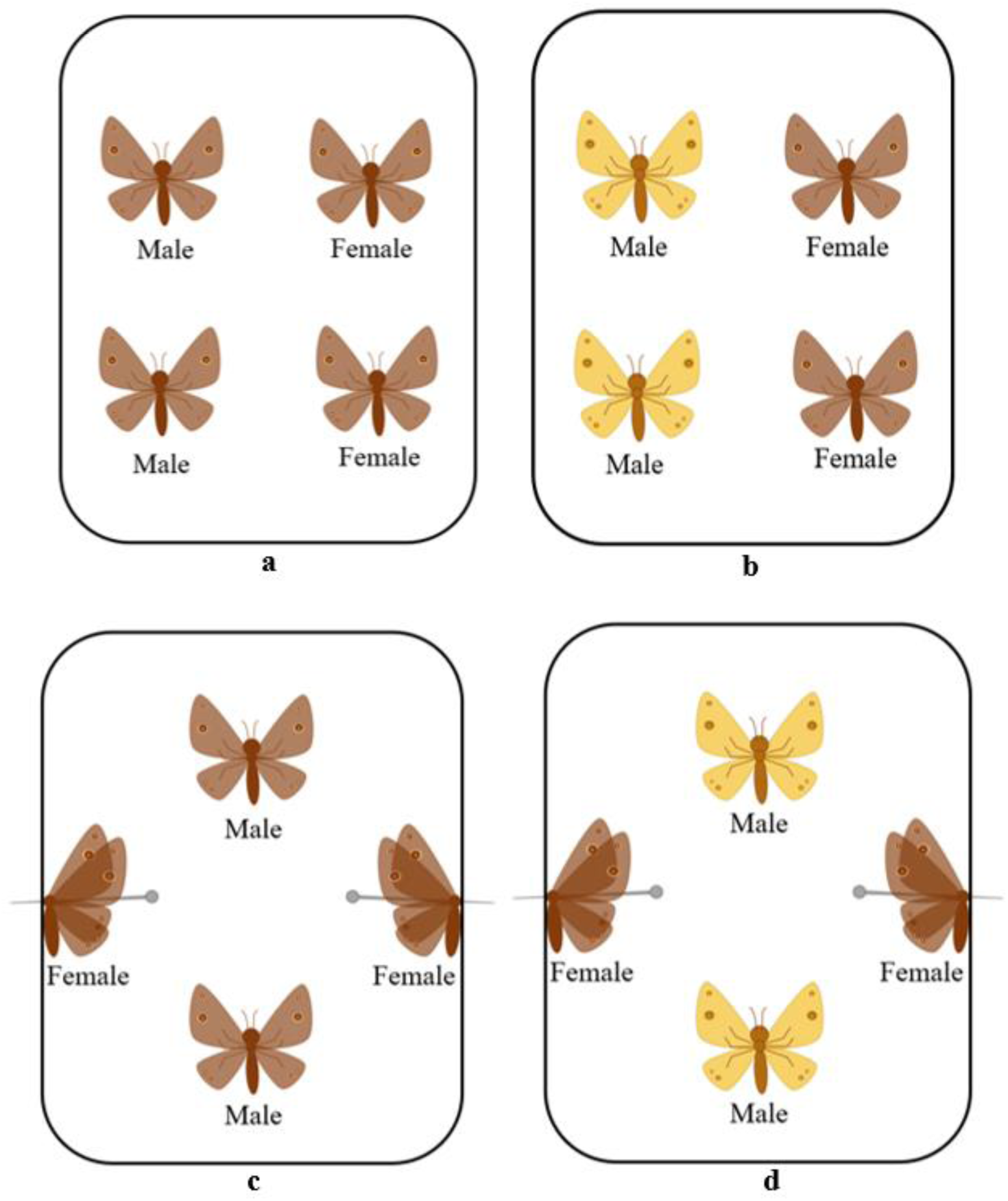
Experimental design of the live **(a-b)** and decapitation **(c-d)** experiments. Two males and two females (live or decapitated) were put in one cage and left under UV light for one hour of observation. Quantification of courtship begins once the UV light is switched on, until the mating of individual butterflies. **(a)** Two Wt males and two Wt females; **(b)** two Yellow males and two Wt females; **(c)** two Wt males and two decapitated Wt females; **(d)** two Yellow males and two decapitated Wt females. These were done on wet season (WS) and dry season (DS) butterflies.

**Supplementary Methods for Fig. S5 (next page)**

*UV photography and spectrophotometry*

To measure the UV reflectivity of the Yellow mutant and Wt male forewing eyespots on both the ventral and dorsal sides (Fig. S5), WS butterflies were photographed using a Nikon D7100 digital camera with a Jenoptik CoastalOpt 105 mm UV – Vis crystal lens under sunlight. Visible light was captured through a Baader UV/IR Cut Filter (transmits 400 to 680 nm) and UV images were taken through a Baader U-Venus Filter (transmits 320 to 380 nm). The camera settings were ISO100 and a shutter speed of 1/320 seconds for visible light and 10 seconds for UV light.

Scale reflectance was measured at the two forewing eyespots using a gonio-spectrophotometer and the accompanying program, OceanView 1.6.7 (Ocean Optics). Each measurement was taken with the axis of the illuminating and detecting fibre directed at a 20° angle to the plane of the wing at a using a deuterium-halogen tungsten lamp (DH-2000, Ocean Optics) as a standardized light source and calibrated using a white Ocean Optics WS-1 reflectance standard. from the right forewing. A total of three replicates were done for each type and sex.

**Fig. S5.**
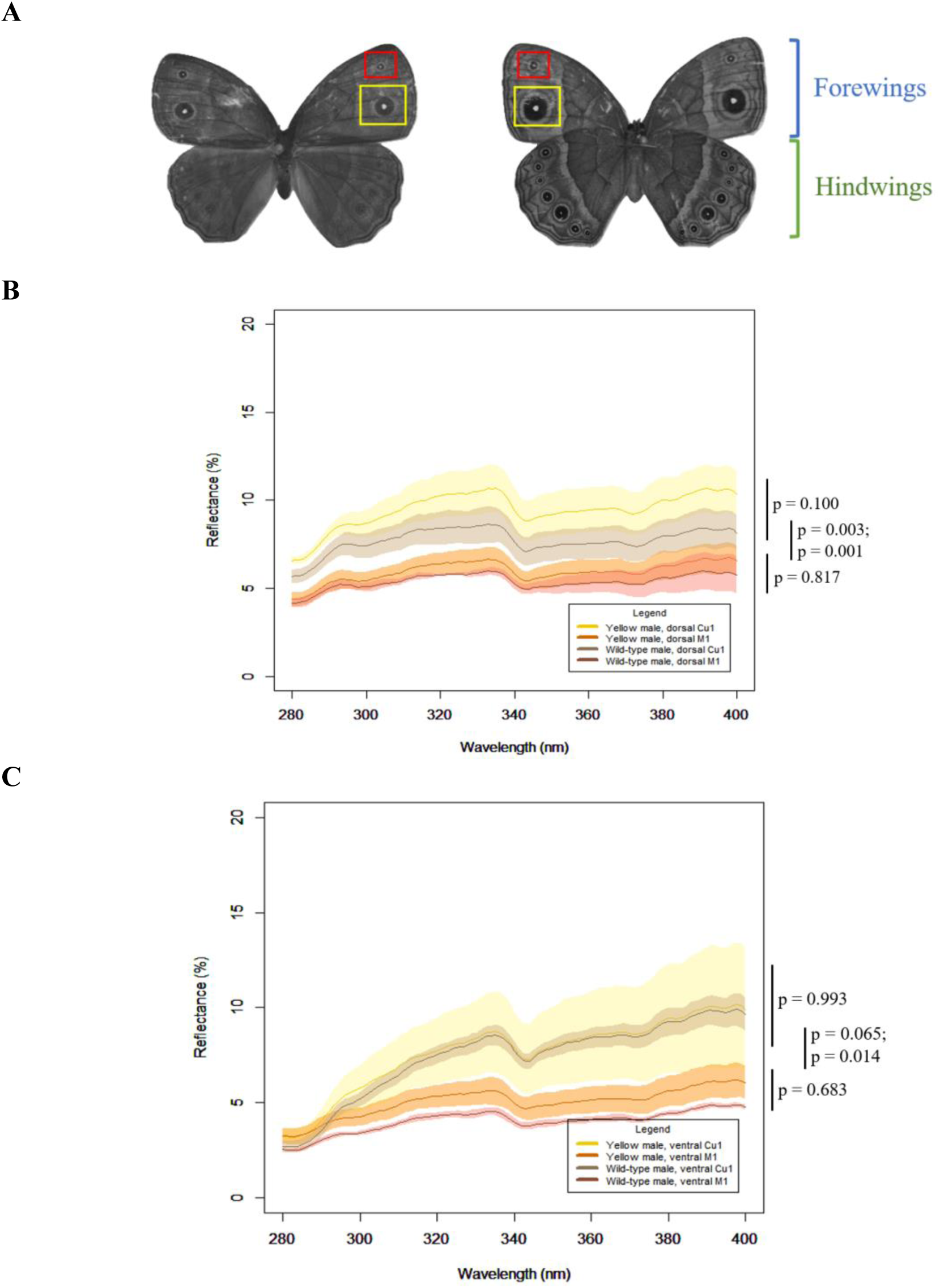
WS Yellow mutant and WT males have similar UV reflectivity of both forewing eyespots on both sides of the wing. **(A)** shows both the dorsal (left) and ventral (right) sides of a Wt butterfly, M1 and Cu1 eyespots are indicated by red and yellow squares respectively. Graphs show the mean smoothed reflectance spectra (±standard deviation, n=3) of WS Yellow mutant and Wt butterflies (280 – 400 nm). The legend for each graph is shown on the bottom right: yellow – Yellow mutant Cu1, orange – Yellow mutant M1, brown – Wt Cu1, red – Wt M1. **(B)** Male Yellow and Wt *B. anynana,* dorsal Cu1 and M1; **(C)** Male Yellow and Wt *B. anynana*, ventral Cu1 and M1.

**Fig. S6.**
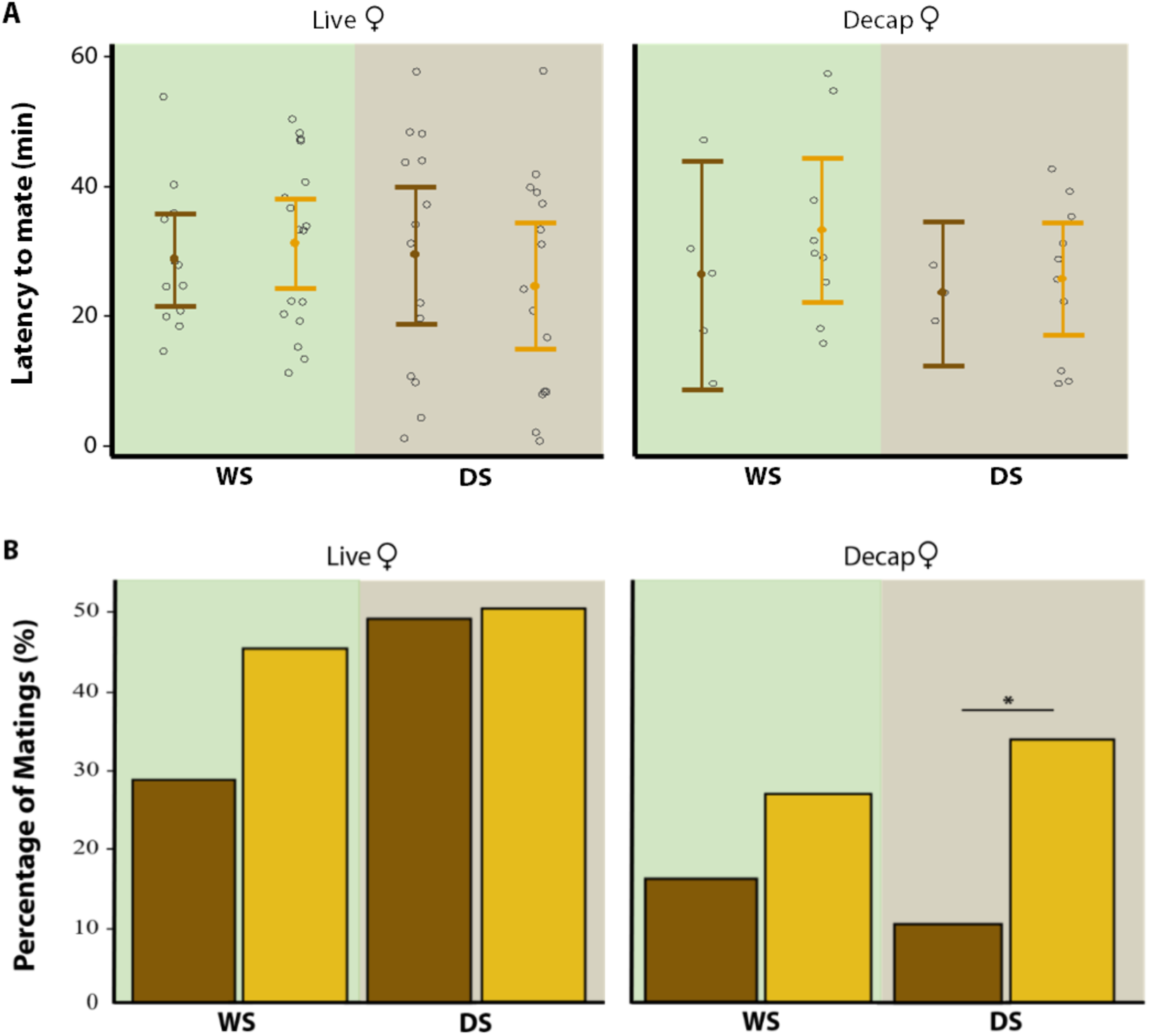
Courtship behavior of Wt and Yellow males, for wet (WS) and dry (DS) seasonal forms, in both live and decapitation assays. **(A)** Latency to mate and **(B)** Percentage of matings were quantified among mated males only. Vertical bars represent 95% confidence intervals. Open circles (°) are data points. Asterisks (*) indicate significant differences: * p ≤ 0.05, ** p ≤ 0.01, *** p ≤ 0.001. Outliers are not removed as they are true measurements. n(WS-Live-Wt) and n(WS-Live-Yellow) = 38, n(WS-Decap-Wt) and n(WS-Decap-Yellow) = 34, n(DS-Live-Wt), n(DS-Live-Yellow), n(DS-Decap-Wt) and n(DS-Decap-Yellow) = 30.

**Fig S7.**
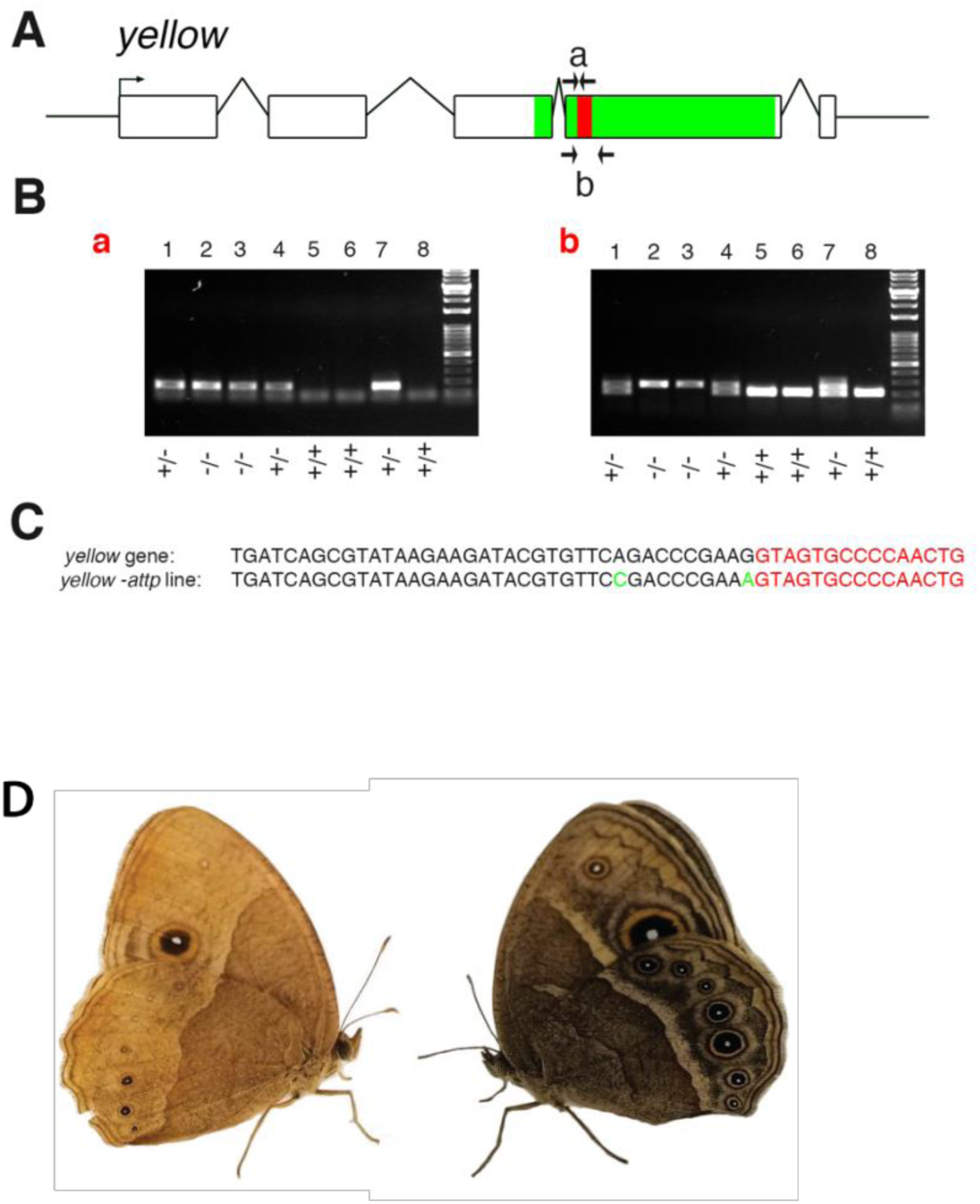
Generation of the Yellow-attP mutant knockout line. **(A)** Insertion of an attP sequence into exon 4 (annotated using LepBase ensemble), the attP insertion is shown in red, the major royal jelly domain is shown in green. **(B)** Gel images of the haemolymph PCR for identifying transgenic individuals. Gel a shows the PCR products using primers designed to the attP insert. Gel b shows the PCR products using primers designed to the *yellow* coding sequence spanning the attP insertion which reveals which animals are heterozygous or homozygous based on the number and size of the PCR band. Individual no. 1 shows 2 bands thus has both the Wt and attP genome. Individual no. 2 shows only 1 band which is slightly larger meaning it has the attP genome only. Individual no. 5 shows only a single small band thus has the Wt genome only. **(C)** Alignment of the yellow sequence with the corresponding region from the yellow-attP line. **(D)** Phenotype of the Yellow mutant knockout butterflies (left) and Wt (right).

